# Neuronal connectivity, behavioral, and transcriptional alterations associated with the loss of MARK2

**DOI:** 10.1101/2023.12.05.569759

**Authors:** Hanna O. Caiola, Qian Wu, Shaili Soni, Xue-Feng Wang, Kevin Monahan, Zhiping P. Pang, George C. Wagner, Huaye Zhang

**Affiliations:** Department of Neuroscience and Cell Biology, Rutgers Robert Wood Johnson Medical School, Piscataway, New Jersey, United States of America; Child Health Institute of New Jersey, New Brunswick, New Jersey, United States of America; Department of Molecular Biology and Biochemistry, Rutgers University, Piscataway, New Jersey, United States of America; Department of Psychology, Rutgers University, Piscataway, New Jersey, United States of America

**Author notes:** These authors contributed equally (Hanna O. Caiola, Qian Wu).

## Abstract

Neuronal connectivity is essential for adaptive brain responses and can be modulated by dendritic spine plasticity and the intrinsic excitability of individual neurons. Dysregulation of these processes can lead to aberrant neuronal activity, which has been associated with numerous neurological disorders including autism, epilepsy, and Alzheimer’s disease. Nonetheless, the molecular mechanisms underlying aberrant neuronal connectivity remains unclear. We previously found that the serine/threonine kinase Microtubule Affinity Regulating Kinase 2 (MARK2), also known as Partitioning Defective 1b (Par1b), is important for the formation of dendritic spines *in vitro.* However, despite its genetic association with several neurological disorders, the *in vivo* impact of MARK2 on neuronal connectivity and cognitive functions remains unclear. Here, we demonstrate that loss of MARK2 *in vivo* results in changes to dendritic spine morphology, which in turn leads to a decrease in excitatory synaptic transmission. Additionally, loss of MARK2 produces substantial impairments in learning and memory, anxiety, and social behavior. Notably, MARK2 deficiency results in heightened seizure susceptibility. Consistent with this observation, RNAseq analysis reveals transcriptional changes in genes regulating synaptic transmission and ion homeostasis. These findings underscore the *in vivo* role of MARK2 in governing synaptic connectivity, cognitive functions, and seizure susceptibility.

## Introduction

The precise establishment and refinement of neuronal connectivity are critical for the brain to adapt and respond to its environment. In the human brain, neurons communicate through trillions of synaptic connections, leading to the formation of complex neuronal ensembles that mediate higher order cognitive functions such as learning and memory. Most of the excitatory synapses are formed on dendritic spines, which are small protrusions on dendrites that play key roles in modulating connectivity in the brain (1). Neural network connectivity is also intricately regulated by the intrinsic excitability of neurons. Increased excitability can facilitate the activation of subthreshold synaptic connections, which in turn may strengthen the connection and support memory formation (2). However, maladaptive changes in neuronal connectivity and excitability can lead to neurological disorders such as autism spectrum disorders (ASD) (3,4), epilepsy (5), and Alzheimer’s disease (AD) (6–8). Nonetheless, the molecular mechanisms underlying neuronal connectivity and how dysregulation of this process may lead to neurological disorders remain unclear.

We previously found that the Microtubule Affinity Regulating Kinase 2 (MARK2), also known as Partitioning Defective 1b (Par1b), is important for the formation of dendritic spines in primary hippocampal neurons (9). MARK2 is a serine/threonine kinase that has key roles in cell polarity establishment and microtubule dynamics in many different cellular contexts (10–14). MARK2 is expressed at high levels in the brain and has been implicated in several neurological disorders such as ASD (15–17), bipolar disorder (18), schizophrenia (19), major depressive disorders (20), and AD (21–23). Previous *in vitro* studies show MARK2 plays a role in establishing neuronal polarity and regulating dendritic spine formation through modulating microtubule dynamics and the synaptic scaffold (9,23–27). Moreover, MARK2 activity is required for proper neuronal migration (28). Nonetheless, it remains unclear how loss of MARK2 affects neuronal connectivity and cognitive functions *in vivo*.

Here, we show that loss of MARK2 *in vivo* results in increased thin dendritic spines at the expense of mushroom spines, with a concomitant reduction in excitatory transmission in the hippocampus. Moreover, loss of MARK2 produces profound behavioral defects including impaired learning and memory, reduced anxiety-like behaviors, and impaired social behavior. Surprisingly, we also found that loss of MARK2 results in increased seizure susceptibility. To understand molecular pathways that may contribute to these phenotypes, we performed RNAseq in MARK2-/- hippocampi and found that pathways involved with ion homeostasis and synaptic transmission are primarily dysregulated. Together, our results establish the *in vivo* roles of MARK2 in regulating synaptic connectivity and behavior and provide insight into the molecular mechanisms underlying the function of MARK2 in the brain.

## Results

### Expression of MARK family proteins in MARK2 knockout mice

To examine the *in vivo* functions of MARK2 in neuronal connectivity and behavior, we used *Mark2* knockout (KO) mice (B6.129X1-*Mark2*^tm1Hpw^/J) (29). In mammals, there are four Par1/MARK members (13). To determine whether knockout of Par1b/MARK2 *in vivo* causes compensatory expression of other Par1/MARK family members in the brain, total protein was extracted from forebrain tissue of 5- to 8-week-old MARK2 KO mice and expression level of different Par1/MARK family members were examined by Western blot. Expression of Par1b/MARK2 was partially decreased in MARK2+/- and almost completely diminished in MARK2-/- mice as expected **(Fig 1A** and B).

**Fig. 1.**
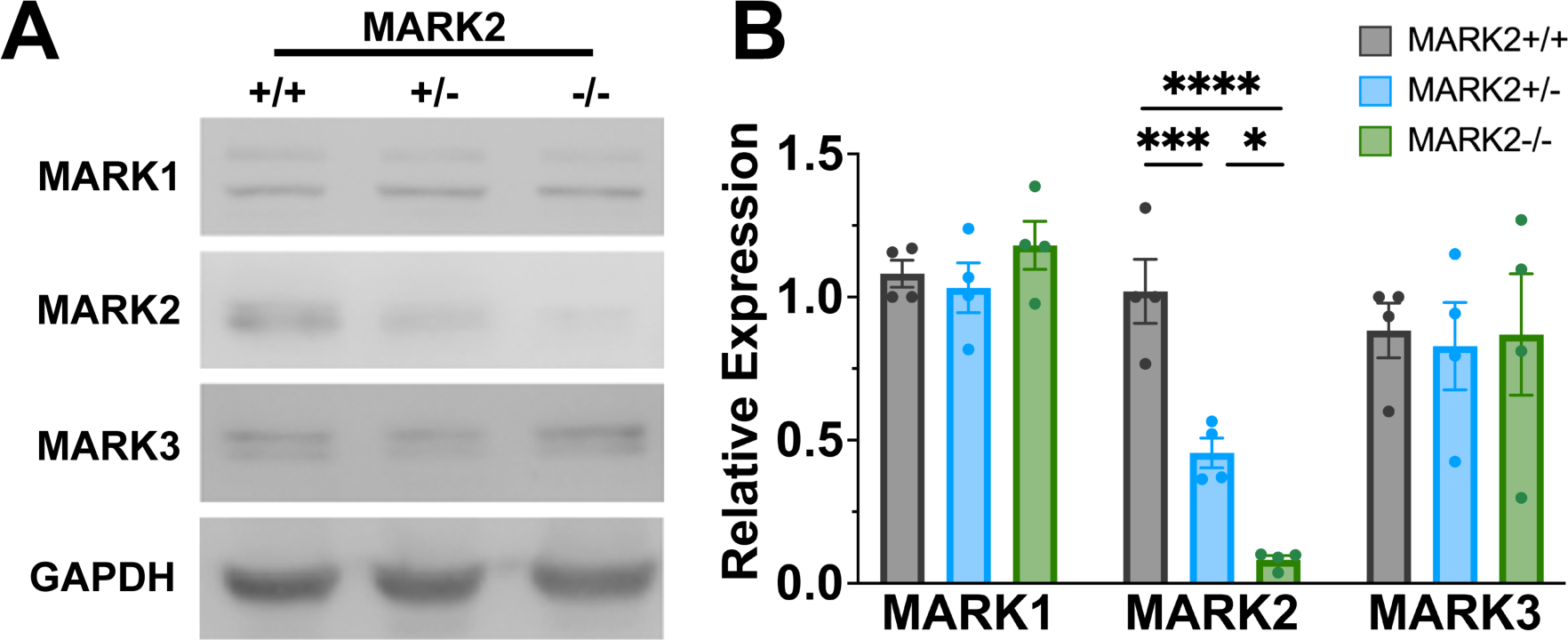
Expression of MARK family proteins in MARK2 knockout mice **(A)** Representative blot of MARK1, MARK2, and MARK3 protein expression in MARK2+/+ (n=4), MARK2+/- (n=4), and MARK2-/- (n=4) normalized to GAPDH. **(B)** Quantification of MARK protein expression from Western blot in (A). One-way ANOVA was performed followed by Tukey’s *post hoc* test: MARK1 (F(2,9)=1.107, p=0.3397), MARK2 (F(2,9)=43.39, p<0.0001), and MARK3 (F(2,9)=0.03143, p=0.9692). *p<0.0332, **p<0.0021, ***P<0.0002, **** p<0.0001. Plot shows mean +/- SEM.

Expression of Par1c/MARK1 or Par1a/MARK3 was not significantly altered in either the MARK2 +/- or -/- mice **(Fig 1).** Par1d/MARK4 is expressed at very low levels in the brain and therefore could not be reliably detected with Western blot. Altogether, these results confirm that MARK2 is decreased as expected in MARK2+/- and MARK2-/- animals while other MARK family members are not affected.

### Loss of MARK2 leads to defects in dendritic spine morphogenesis and synaptic transmission in the hippocampus

Dendritic spines are sites of excitatory input in the nervous system. Changes to dendritic spine density, size, and shape have been linked to numerous neurological disorders (30,31). The size and shape of spines range from immature filipodia, which are long, thin spines to mature mushroom-shaped spines, which are shorter and wider (32). Furthermore, mushroom-shaped spines generally contain larger postsynaptic density with an increase in AMPA receptors (33,34). Therefore, mushroom spines are considered more mature than longer and thinner spines due to their increased synaptic strength (35).

Previously, we and others have shown that Par1/MARK is necessary for spine morphogenesis in cultured hippocampal neurons (9,27). To explore whether MARK2 plays a role in spine morphogenesis *in vivo*, we crossed MARK2 KO mice with Thy1-YFP mice to create MARK2 KO/YFP-H mice (36), in which the layer V pyramidal neurons in the cortex and pyramidal neurons in the hippocampus are positive for YFP. Dendritic spines of pyramidal neurons on the secondary apical dendrites in the *strata radiatum* layer were measured for density, length, width, and morphological classification **(Fig 2A)**. We found no differences across genotypes for length, width **(Fig 2B)**, or overall spine density **(Fig 2C)**. Interestingly, we found that MARK2+/- and MARK2-/- mice have an increased number of thin spines and reduced number of mushroom spines **(Fig 2D)**. These results suggest that while the overall density of dendritic spines is not altered, there is a shift towards increased immature spines in MARK2+/- and MARK2-/- mice.

**Fig. 2.**
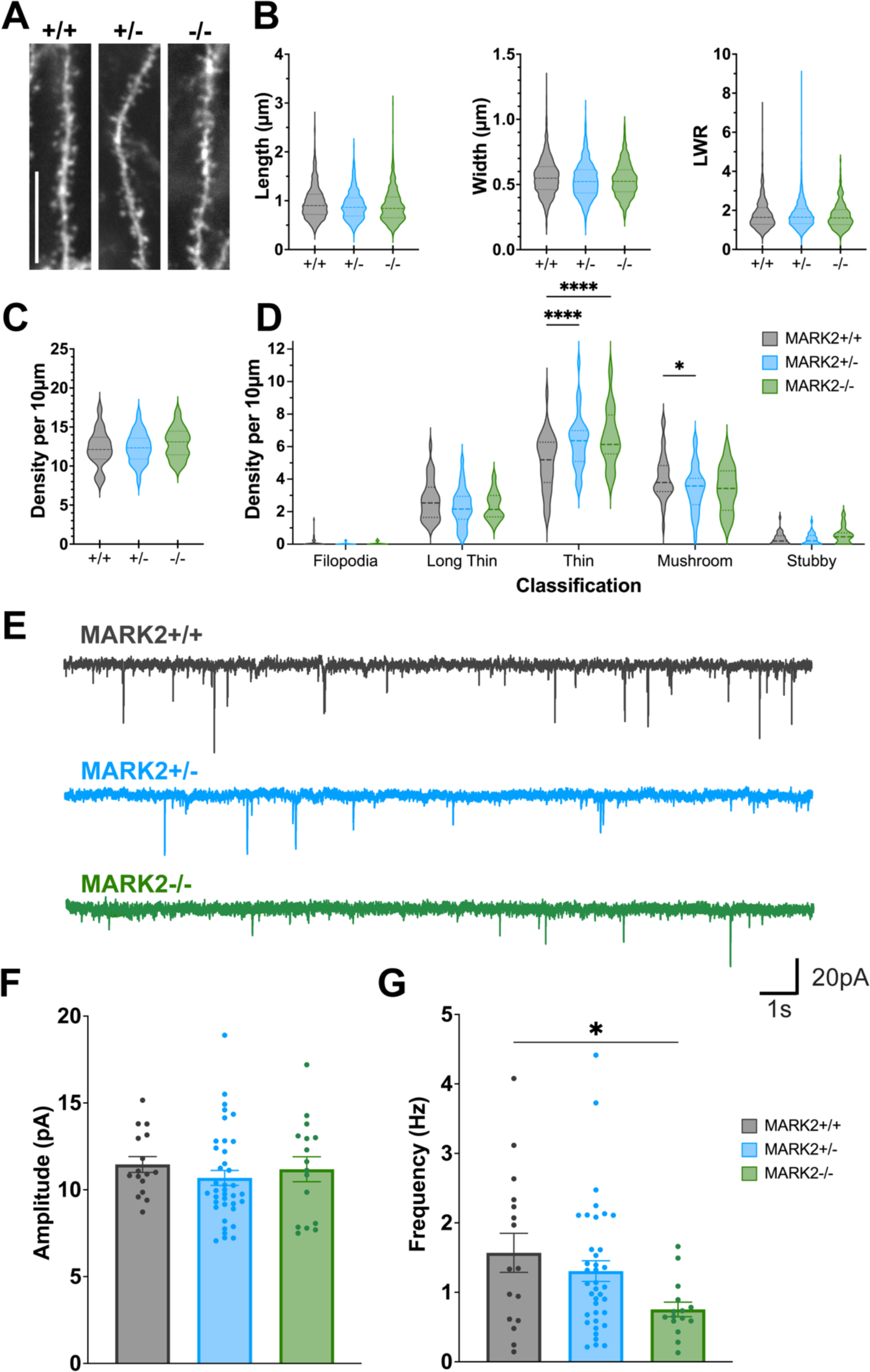
Loss of MARK2 leads to defects in dendritic spine morphogenesis and synaptic transmission in the hippocampus. **(A)** Representative images of secondary dendrites of pyramidal neurons in *strata radiatum* layer of the hippocampus from MARK2+/+ (n=4 mice, 35 dendrites, 1858 spines), MARK2+/- (n=4 mice, 31 dendrites, 1642 spines), and MARK2-/- (n=2 mice, 16 dendrites, 911 spines) mice. Scale bar: 10µm. **(B)** No changes in spine length (F(2,7)=2.242, p=0.1768), spine head width (F(2,7)=2.038, p=0.2006), and length-to-width ratio (LWR; F(2,7)=0.6789) were detected between genotypes by nested one-way ANOVA . **(C)** No changes in overall spine density were observed between genotypes by nested one-way ANOVA (F(2,7)=0.1386, p=0.8729). **(D)** Spines were morphologically classified sequentially using the following criteria: filipodia (length>2µm), mushroom (head width>0.6µm), long thin (length>1µm), thin (LWR>1µm), or stubby(LWR<1µm). MARK2+/- and MARK2-/- dendrites had fewer mushroom spines and increased thin spines. Two-way ANOVA followed by Tukey’s *post hoc*. **(E)** Representative mEPSC traces from hippocampal neurons of MARK2+/+, MARK2+/-, and MARK2-/- mice. **(F)** mEPSC amplitude compared between MARK2+/+ (n=16), MARK2+/- (n=39), and MARK2-/- (n=16) neurons. Kruskal Wallis ANOVA (stat=2.391, p=0.3025). **(G)** mEPSC frequency between MARK2+/+, MARK2+/-, and MARK2-/- neurons. Brown-Forsythe ANOVA (F*(2, 32.10)=3.678, p=0.0364) followed by Dunnett’s T3 *post hoc* test. *p<0.0332, **p<0.0021, ***P<0.0002, **** p<0.0001. Mean +/- SEM.

Our finding of increased immature spines in MARK2+/- and MARK2-/- mice suggests potential changes in synaptic transmission since immature spines are less functional than their mature counterparts. Thus, we aimed to understand if there were any functional defects at the synaptic level. Miniature excitatory postsynaptic currents (mEPSCs) in hippocampal pyramidal neurons were measured by whole-cell voltage clamp in acute brain slices **(Fig 2E)**. We found no changes in mEPSC amplitude **(Fig 2F)**; however, a significant decrease in mEPSC frequency was observed in MARK2-/- mice when compared with MARK2+/+ mice **(Fig 2G)**. Together, these results suggest that the MARK2 KO mice exhibit a net reduction in functional synapses as measured by mEPSC frequency. This is in line with the increased number of thin spines and reduction of mushroom-shaped spines observed in the MARK2 KO mice.

### Loss of MARK2 results in deficits in hippocampal-dependent learning and memory tasks

Changes to dendritic spines and synaptic transmission are often correlated with changes at the behavioral level. In particular, proper regulation of dendritic spines and synaptic transmission is critical in learning and memory tasks (37,38). Therefore, we aimed to understand whether MARK2+/- and MARK2-/- mice exhibit deficits in learning and memory. First, we used the hippocampal-dependent learning task Morris water maze (MWM). MARK2+/+, MARK2+/-, and MARK2-/- mice were trained for 2 days with the platform visible and then for an additional 4 days with the platform hidden under the surface of the water with each training day consisting of 4 trials **(Fig 3A)**. During training days 1 and 2 in which the platform was visible, all mice successfully found the platform; however, by the last trial on Day 1, MARK2-/- mice showed an increased latency to platform compared to MARK2+/+ and MARK2+/- mice **(S1 Fig)**. Once the platform was hidden on training Day 3 through Day 6, MARK2-/- mice consistently exhibited increased latency to platform compared to MARK2+/+ mice **(Fig 3B)**.

**Fig. 3.**
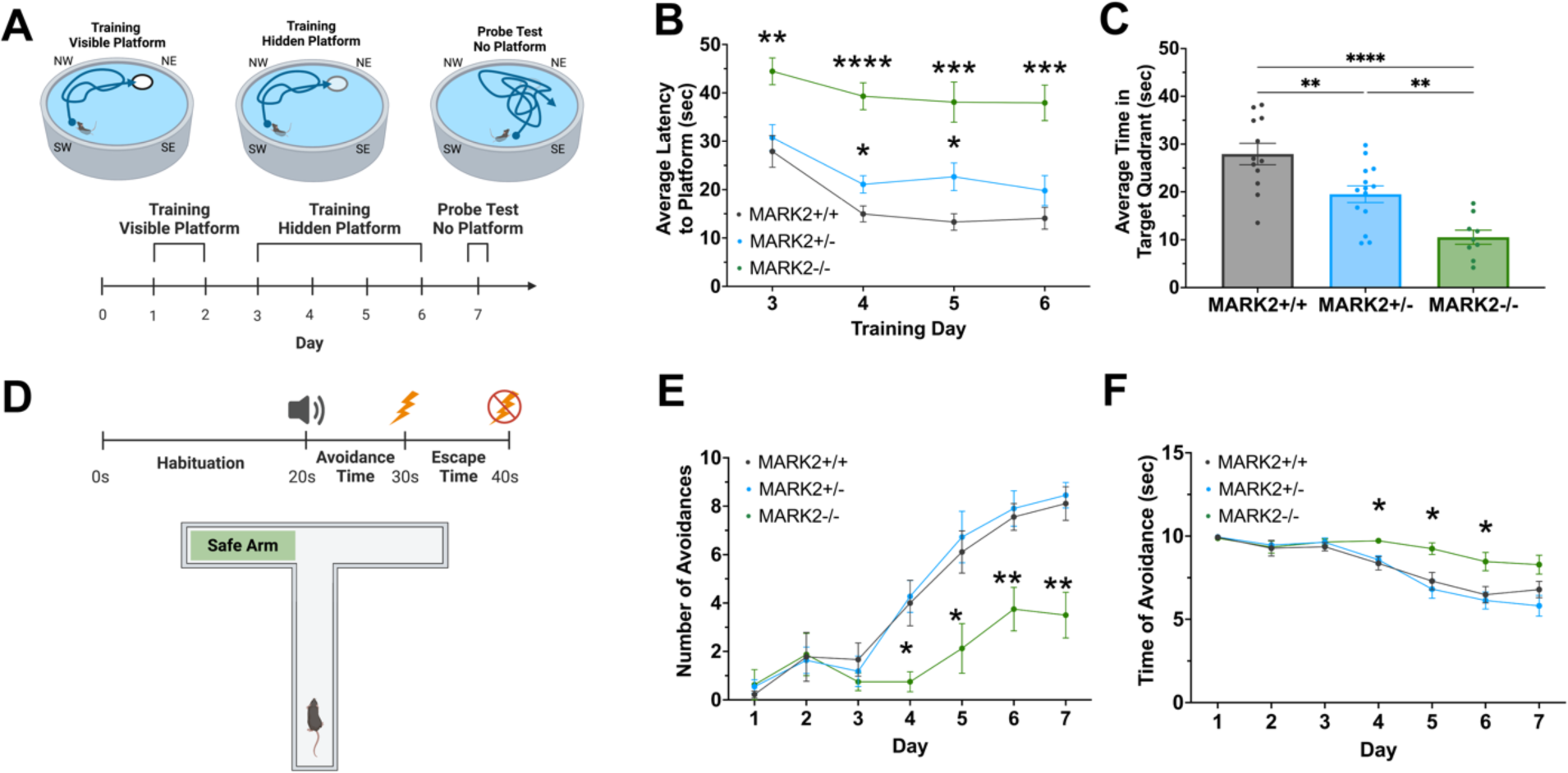
Loss of MARK2 results in deficits in hippocampal-dependent learning and memory tasks. **(A)** Experimental timeline of MWM training. The platform was visible during Days 1 and 2 of training and then hidden during Days 3 through 6. The platform was removed completely during the probe test. Each animal underwent 4 trials per day. **(B)** Average latency to platform in seconds on training days 3 through 6 (hidden platform) between MARK2+/+ (n=12), MARK2+/- (n=14), and MARK2-/- (n=9) mice. Two-way repeated measures ANOVA, with Dunnett’s *post hoc* test. **(C)** Performance of MARK2+/+ (n=12), MARK2+/- (n=14), and MARK2-/- (n=9) mice in probe test on Day 7. Ordinary one-way ANOVA (F(2,32)=18.50, p<0.0001) followed by Tukey’s *post hoc*. **(D)** Experimental timeline of each active avoidance task trial. Mice were placed into a T-maze and allowed to habituate. After 20 seconds of habituation, a tone initiated the trial and the door to the safe arm opened. After 10 seconds, a 0.8 mA foot shock was administered for an additional 10 seconds. The mice were considered to have successfully avoided the shock if they reached the safe arm prior to shock onset. Each mouse underwent 10 trials per day. **(E)** Number of times MARK2+/+ (n=10), MARK2+/- (n=10), and MARK2-/- (n=8) mice avoided the shock. Two-way repeated measures ANOVA followed by Dunnett’s *post hoc* test. **(F)** Time it took for MARK2+/+, MARK2+/-, and MARK2-/- mice to reach the safe arm after the tone was played. Two-way repeated measures ANOVA followed by Dunnett’s *post hoc* test. *p<0.0332, **p<0.0021, ***P<0.0002, **** p<0.0001. Mean +/- SEM.

MARK2+/- mice exhibited increased latency to platform compared to controls only on Days 4 and 5 **(Fig 3B)**. During the probe test on Day 7 in which the platform was removed from the tank, MARK2-/- mice spent significantly less time in the target quadrant compared to the MARK2+/+ and MARK2+/- mice **(Fig 3C)**. MARK2+/- mice also performed significantly worse in the probe test compared to MARK2+/+ mice **(Fig 3C)**.

Next, we performed another hippocampal-dependent task, the T-maze active avoidance test **(Fig 3D)**. The mice were subject to 10 trials per day. Each trial consisted of an initial interval of 20 seconds after which a tone was played to initiate the trial. The mice had 10 seconds to reach the escape arm, which would avoid a 10 second 0.8 mA foot shock. If the mice failed to avoid the shock, they could escape after shock onset by moving to the escape arm. Mice were considered to have learned the task if they successfully paired the conditioned stimulus (tone) with the unconditioned stimulus (shock) and exhibited avoidance behavior.

Given the impaired performance of the MARK2-/- mice in the MWM, we hypothesized that these mice would have increased avoidance times and reduced number of avoidances compared to MARK2+/+ and MARK2+/- mice. We found that MARK2+/+ and MARK2+/- mice successfully learned to associate the tone and shock by Day 4 as evidenced by and increased number of avoidances **(Fig 3E)** and decreased avoidance times **(Fig 3F)**, which continued to trend in the same respective directions through Day 7. Interestingly, the MARK2-/- mice seemed to eventually learn to associate the tone and shock, but at a much slower rate. The number of avoidances exhibited by the MARK2-/- mice began to increase on Day 5 and continued through Day 6; however, the total number of avoidances was significantly lower from Day 4 through Day 7 compared to MARK2+/+ and MARK2+/- **(Fig 3E)**. Additionally, even though the number of avoidances moderately increased each day, it took the MARK2-/- mice significantly longer to avoid the shock **(Fig 3F)**. To ensure that the poor performance of the MARK2-/- mice were not a result of decreased motor or locomotor activity, the mice were also subjected to rotarod and locomotor tests in which we found no differences between genotypes **(S2 Fig)**. In sum, these experiments indicate that MARK2 is necessary for proper learning and memory formation.

### MARK2-/- mice show reduced sociability and reduced anxiety-like behavior

Altered dendritic spine morphology and synaptic transmission have been shown to coincide with deficits in social interaction and altered anxiety levels (39–41). In addition, MARK2 is associated with neurological disorders that often exhibit social deficits and anxiety (15–17,20–23,42). Therefore, we wanted to explore whether MARK2+/- and MARK2-/- mice exhibit deficits in these behaviors. Indeed, we found that MARK2-/- mice exhibit impaired social behavior in the three-chamber social interaction test. Subject mice were placed into the center chamber with an empty cylinder in the end chamber (control) and a cylinder containing a sex- and age-matched heterozygous mouse target in the other end chamber (target) **(Fig 4A)**. To measure social interaction, the number of times the subject mouse contacted each cylinder and the total amount of time the subject mouse spent interacting with each cylinder were recorded. Intriguingly, we found that MARK2-/- mice exhibited fewer target cup touches compared to the MARK2+/+ mice **(Fig 4B)**. In addition, the MARK2-/- mice trended towards a decrease in the amount of time spent interacting with the target compared to MARK2+/+ controls; however, this did not reach significance **(Fig 4C)**. Together, these results suggest that MARK2-/- mice exhibit impaired social interactions and that MARK2 plays an important role in regulating social behavior.

**Fig. 4.**
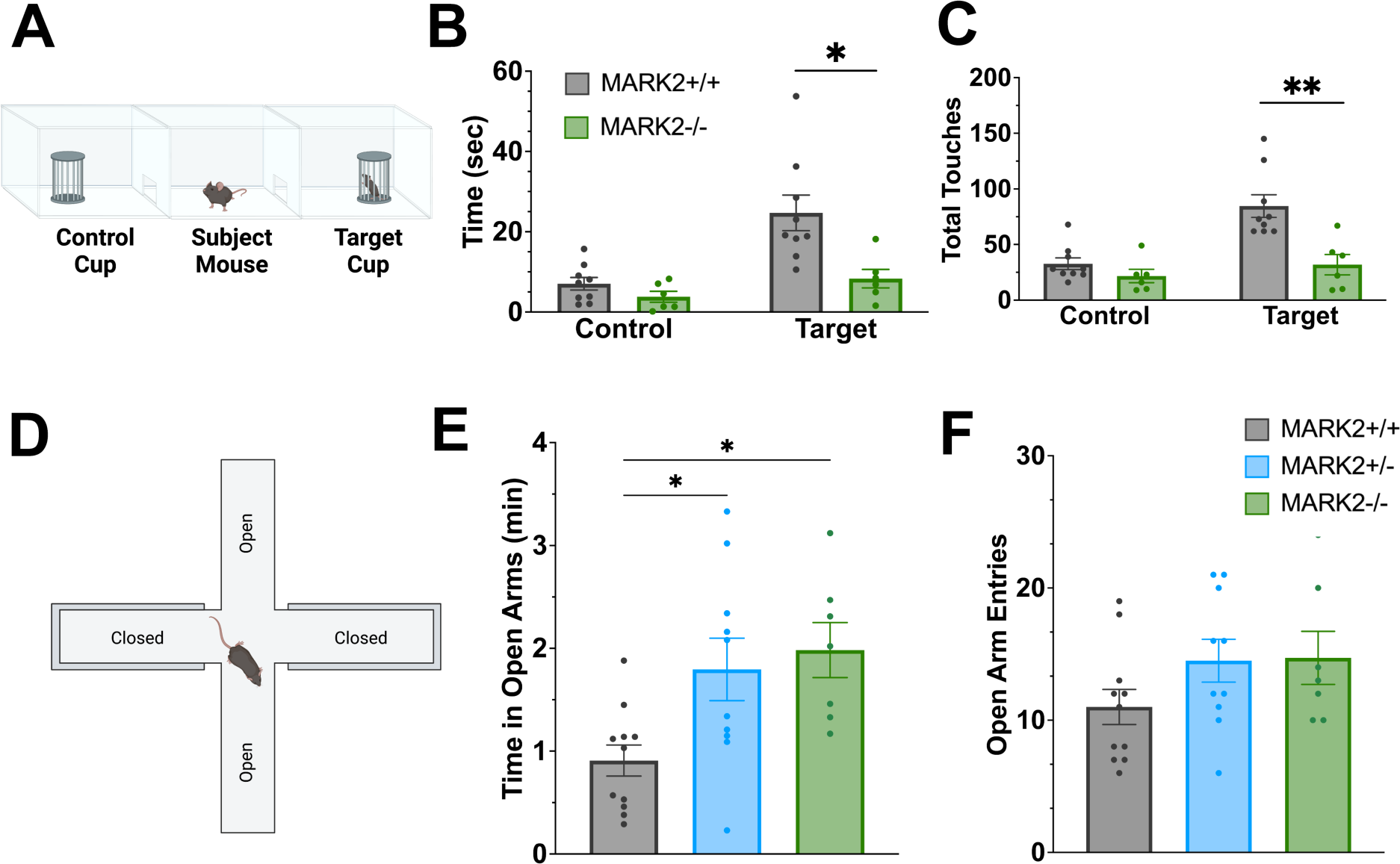
MARK2-/- mice show reduced sociability and anxiety-like behavior. **(A)** Diagram of three-chamber sociability test. Mice were habituated to the chamber for 10 minutes. After habituation, an age- and sex-matched MARK2+/- mouse was placed in the target cup and the subject mouse was placed back into the center chamber. The control cup was left empty. **(B)** Total amount of time MARK2+/+ (n=9) and MARK2-/- (n=6) subject mice interacted with the control and target cups. Unpaired t-test was used to test for differences between MARK2+/+ vs. MARK2-/- for control (t=1.451, p=0.1704) and target cups (t=2.822, p=0.0144). **(C)** Total number of times MARK2+/+ and MARK2-/- subject mice touched the control and target cups with one or both paws. Mann Whitney U-test was used to test for differences between MARK2+/+ vs. MARK2-/- for control (U=11.50, p=0.0692) and target cups (U=3, p=0.0028). **(D)** Diagram of the elevated plus maze (EPM). Each mouse was placed in the center square and observed for 10 minutes. **(E)** The total amount of time MARK2+/+ (n=13), MARK2+/- (n=12) and MARK2-/- (n=8) spend in the open arms. One-way ANOVA (F(2, 25)=5.743, p=0.0088) followed by Dunnett’s *post hoc* test. **(F)** The total number of open arm entries of MARK2+/+, MARK2+/-, MARK2-/- mice. An entry was defined as when all four paws entered the open arm. One-way ANOVA F(2, 25)=1.796, p=0.1868). *p<0.0332, **p<0.0021, ***P<0.0002, **** p<0.0001. Plot shows mean +/- SEM.

Next we wanted to examine whether anxiety was affected in MARK2-/- mice. To test the anxiety level of MARK2-/- mice, we used the elevated plus maze, which measures the anxiety induced by open spaces and height. Mice were placed in the center of the plus-shaped maze, which consisted of two long closed arms and two short open arms **(Fig 4A)**. The number of times and duration of time spent in the open arms was recorded. Surprisingly, we found that MARK2+/- and MARK2-/- mice spent more time in the open arms compared to controls. MARK2-/- spent more time compared to MARK2+/- mice as well **(Fig 4B)**. The number of open arm entries trended towards an increase in MARK2+/- and MARK2-/- mice but did not reach significance **(Fig 4C)**. The increased time spent in open arms with partial and total knockout of MARK2 suggests that MARK2-/- mice exhibit less anxiety.

### Heterozygous deletion of MARK2 results in increased seizure susceptibility

Seizures are often found to be comorbid with neurological disorders such as ASD (3,4,43–46), schizophrenia (47,48), and AD (6–8,49–51) all of which have been genetically linked to MARK2. To determine whether loss of MARK2 affects seizure susceptibility, we injected MARK2+/+ and MARK2+/- mice with pilocarpine (300 mg/kg) intraperitoneally to induce seizures. To reduce side effects from peripheral cholinergic inputs, mice were also injected with methyl scopolamine (1 mg/kg) 30 minutes prior to pilocarpine injection. Mice were recorded for 2 hours following the pilocarpine injection and seizure behavior was scored by a blinded observer using the Racine scale as modified by Borges et al(52,53). Unexpectedly, we found that partial loss of MARK2 was enough to significantly increase seizure susceptibility **(Fig 5A)**. Furthermore, the mortality rate was three times higher in MARK2+/- mice compared to MARK2+/+ controls **(Fig 5B)**. Since this phenotype was strong in MARK2+/- mice and the mortality rate was significantly elevated compared to MARK2+/+ mice, we did not perform this experiment on MARK2-/- mice as the result would have likely been much more fatal. Altogether, these results suggest that partial loss of MARK2 is sufficient to result in increased seizure susceptibility. This result is surprising given our findings that MARK2+/- and MARK2-/- mice have a shift towards immature dendritic spines and reduction in excitatory transmission. This indicates that the increased seizure susceptibility may be due to molecular pathways independent of dendritic spines and excitatory synaptic transmission.

**Fig. 5.**
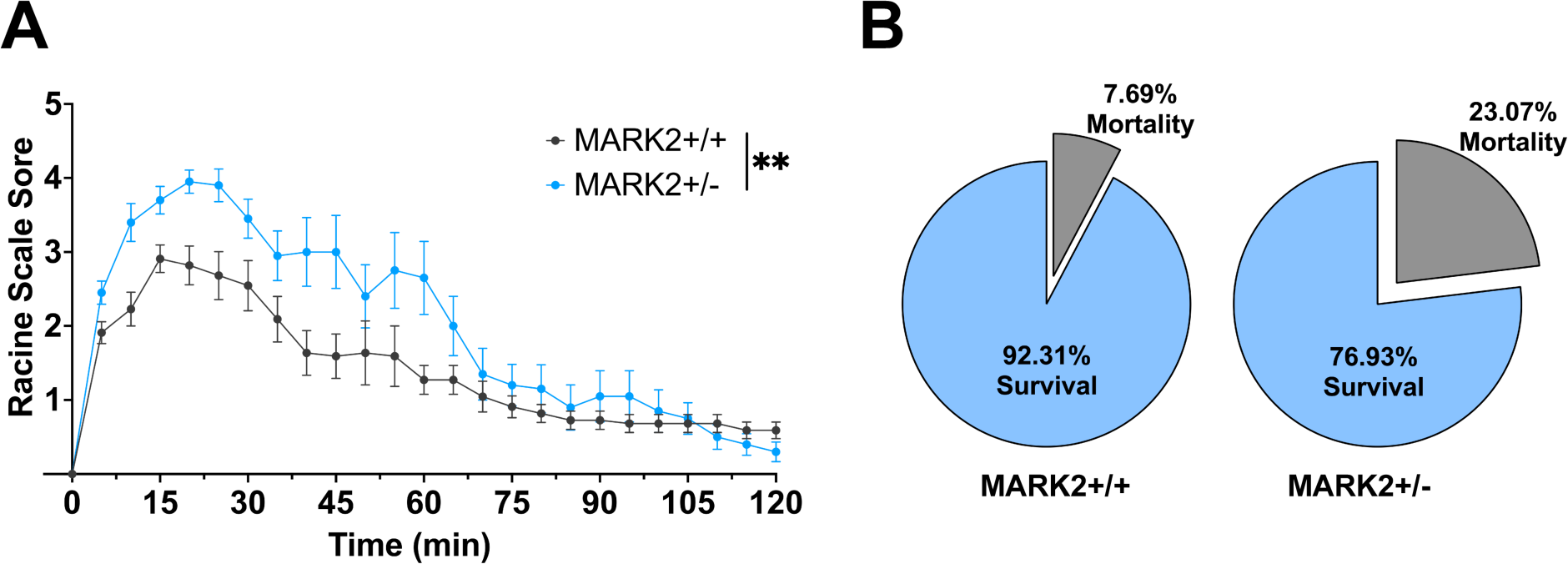
Heterozygous deletion of MARK2 results in increased seizure susceptibility. **(A)** MARK2+/+ (n=13) and MARK2+/- (n=14) mice were injected with pilocarpine (300 mg/kg) to induce seizures. Methyl scopolamine (1 mg/kg) was injected 30 minutes prior to pilocarpine to minimize peripheral cholinergic side effects. Mice were observed for 2 hours and scored on the Racine scale by a blinded experimenter. Two-way repeated measures ANOVA. **(B)** Mortality rate amongst mice used for seizure susceptibility test. *p<0.0332, **p<0.0021, ***P<0.0002, **** p<0.0001. Mean +/- SEM.

### MARK2-/- mice exhibit transcriptional dysregulation of ion channels

To gain insight into the molecular mechanisms that may contribute to the increased seizure susceptibility observed with loss of MARK2, we performed unbiased bulk RNAseq on MARK2+/+ and MARK2-/- hippocampi. We used DESeq2 to perform differential expression analysis and found 522 differentially expressed genes **(Fig 6A,B)**. Among these changes, we observed a significant increase in *Mark2* transcript levels in the MARK2-/- hippocampi; however, we confirmed that exons 2-4 were removed, which should prevent these transcripts from encoding a functional protein **(S4 Fig)**. We observed no significant changes in mRNA levels of *Mark1*, *Mark3* and *Mark4*, which is consistent with our Western blot analyses of MARK proteins **(S4 Fig, Fig1)**. This further confirms that there are no compensatory gene expression changes in other MARK family members.

**Fig. 6.**
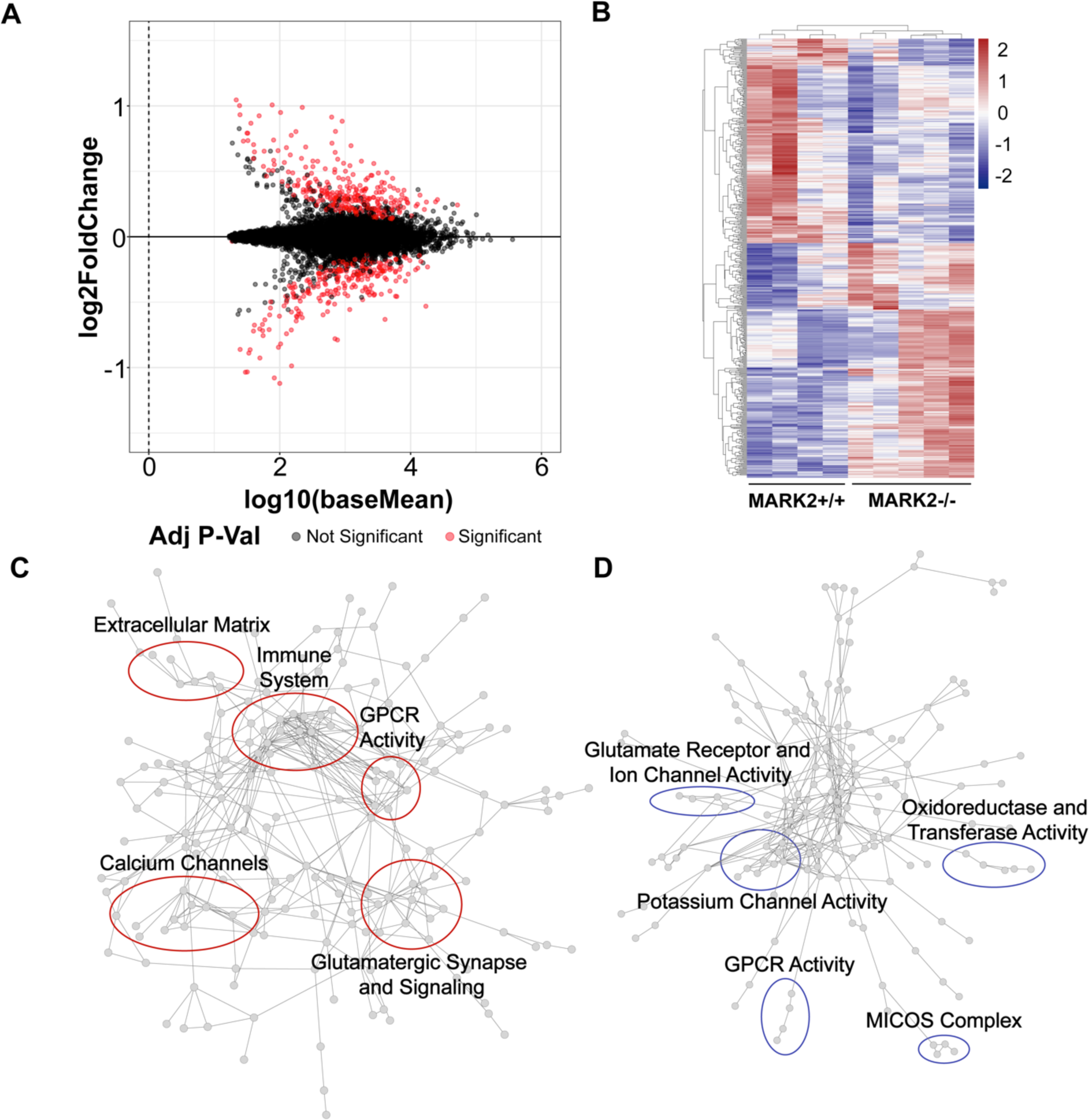
MARK2-/- mice exhibit transcriptional dysregulation of ion channels. **(A)** MA plot of 15,625 genes from DESeq2 analysis of 8-week-old MARK2+/+ (n=4) and MARK2-/- (n=5) mouse hippocampi. Significant genes are shown in red (Padj < 0.05). **(B)** Heatmap of all significant upregulated and downregulated genes identified in DESeq2 analysis. **(C,D)** Networks of upregulated (D) and downregulated (E) genes were made using the STRING database in Cytoscape. The largest subnetwork was selected for subsequent clustering analysis with MCL Cluster AutoAnnotate. The top 5 clusters are circled and labeled with the Gene Ontology (molecular function) term most associated with the genes in each cluster.

To determine which molecular pathways are dysregulated in the MARK2-/- mice, we created a STRING network of upregulated and downregulated genes separately followed by a clustering analysis to find the top 5 up and downregulated clusters. Interestingly, we found that potassium channels and calcium channels are enriched in the upregulated and downregulated genes lists, respectively **(Fig 6C)**. Furthermore, we found that glutamatergic and GPCR activity are enriched in both up and downregulated genes. To further understand the molecular functions enriched in our gene sets independent of their protein-protein interactions, we performed gene ontology analysis in DAVID of up and downregulated genes separately. We found that ion channels, particularly calcium and potassium channels, glutamatergic signaling, and GPCR activity are among the most enriched terms **(Fig 7A,B)**. The presence of ion channels in our STRING and gene ontology analyses could indicate that ion homeostasis is dysregulated with loss of MARK2, which could be a contributing factor to the observed seizure phenotype.

**Fig. 7.**
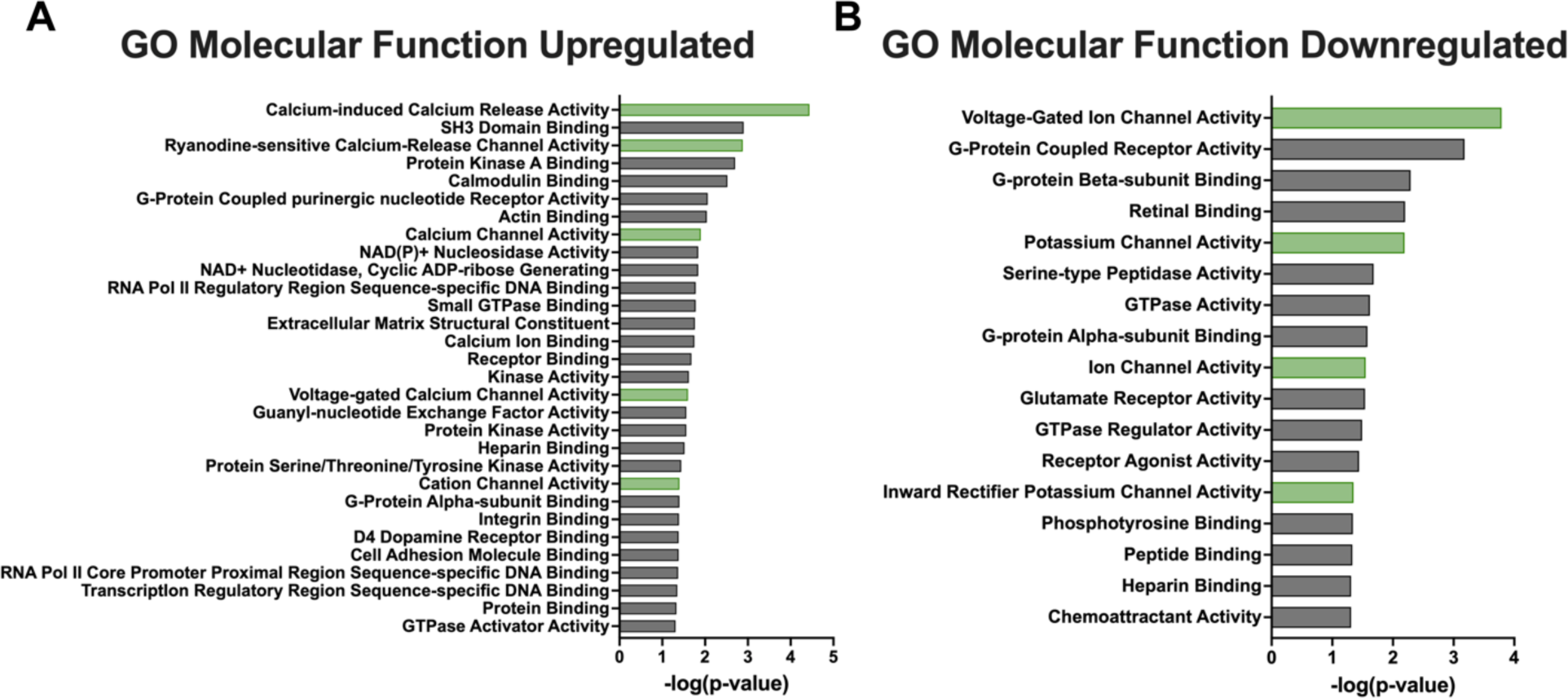
Gene ontology analysis in MARK2-/- mice **(A, B)** Gene Ontology (molecular function) analysis was conducted with DAVID. Terms are sorted in order significance according to the adjusted P-Value. Green bars indicate terms associated with ion channels and ion homeostasis.

## Discussion

In this study, we aimed to understand how MARK2 contributes to synaptic connectivity and cognition *in vivo*. Interestingly, we found that MARK2-/- mice show increased immature dendritic spine formation and decreased mEPSC frequency, indicating a reduction in excitatory synaptic transmission.

The observed spine changes are in line with our previous findings in primary hippocampal neurons and other reports in which MARK2 knockdown *in vitro* results in fewer mature spines and increased filipodia (9,27). To understand the functional impact these changes may have on synaptic transmission in MARK2-/- mice, we also performed electrophysiological recordings and found a decrease in mEPSC frequency, but no changes in amplitude. These results are consistent with another study in which a peptide inhibitor targeting all four MARK family members *in vitro* results in reduced mEPSC frequency, but not amplitude (54).

We performed comprehensive behavioral analyses on MARK2-/- mice. MARK2 is implicated in various neurological disorders such as ASD (15–17), bipolar disorder (18), schizophrenia (19), major depressive disorders (55), and AD (21–23). Thus, we focused our attention on behavioral phenotypes observed in these disorders, including learning and memory, social behavior, and anxiety-related behaviors. Interestingly, we observed profound changes in hippocampal-dependent learning and memory as well as social behaviors in the MARK2-/- mice, which is consistent with the performance of mice from another MARK2 knockout mouse line (EMK1-knockout by gene trapping) (56). Curiously, MARK2-/- mice show increased exploration of the open arm in the elevated plus maze test, indicating decreased anxiety-like behaviors. This is the opposite of the phenotype observed in ASD subjects and animal models, which typically show increased anxiety-like behaviors (57–60). However, increased open arm exploration has been observed in several AD mouse models (61). It will be important in the future to examine how MARK2 activity changes in different neurological disorders such as ASD and AD. Furthermore, conditional knockout mice with brain region specific deletion of MARK2 will provide more insight into the role of MARK2 in different brain areas in controlling various behaviors.

We also examined seizure susceptibility in the MARK2-/- mice, as seizures are often comorbid with ASD, schizophrenia and AD. Remarkably, even heterozygotic deletion of MARK2 results in a significant increase in seizure susceptibility, indicating altered network excitability. As a reduction in excitatory synaptic transmission is typically not expected to coincide with increased seizure susceptibility, we performed an unbiased RNAseq analysis to explore additional molecular pathways affected as a result of MARK2 knockout. We found significant dysregulation in genes associated with synaptic transmission and ion homeostasis. In particular, calcium and potassium channel genes were found to have increased and decreased expression, respectively, in MARK2-/- mice. This suggests that a change in the intrinsic membrane excitability due to disrupted ion homeostasis may underlie the seizure susceptibility in the MARK2-/- mice.

What might be the molecular mechanisms underlying the dysregulation of gene transcription observed in MARK2-/- mice? While MARK2 is not known to be a transcriptional regulator, it has several substrates including CRTC2, CBP, and class IIa histone deacetylases (HDACs), which are transcriptional regulators (62–68). Since phosphorylation of CRTC2 and class IIa HDACs results in their restriction to the cytoplasm, loss of MARK2 would theoretically lead to increased nuclear accumulation of these molecules. Additionally, since MARK2 negatively regulates CBP, loss of MARK2 can lead to increased CBP activity (64). It will be interesting to explore the roles of these transcriptional regulators in modulating synaptic transmission and ion homeostasis in the MARK2-/- mice.

One limitation of the RNAseq experiment is that it is bulk RNAseq performed on whole hippocampal tissue. While the enrichment of synaptic transmission and ion homeostasis genes suggest a neuronal mechanism, many of these genes are also expressed in glial cells. Thus it is possible that dysregulation of gene transcription in glial cells also contributes to some of the observed phenotypes.

Indeed our previous studies found that loss of MARK2 in microglia primes microglia during development and increases their sensitivity to injury (69). It will be of great interest to further examine the cell type specific changes in gene regulation in the MARK2-/- mice.

In conclusion, our study elucidates the *in vivo* functions of MARK2 in regulating neuronal connectivity and behavior. In addition, RNAseq analyses show gene dysregulation in synaptic transmission and ion homeostasis, which provide insight into the underlying molecular mechanisms responsible for the behavioral and seizure phenotypes. Further investigation into the specific molecular pathways and downstream targets of MARK2 will enhance our understanding of the underlying mechanisms and facilitate the development of targeted therapeutic strategies for neurological disorders associated with MARK2 dysregulation.

## Materials and methods

### Animals

All animals used in this study were carried out in accordance with Rutgers-RWJMS Institutional Animal Care and Use Committee protocols. B6.129X1-*Mark2*^tm1Hpw^/J (stock# 009365) and B6.Cg-Tg (Thy1-YFP) HJrs/J (stock# 003782) mice were purchased from Jackson Laboratories. Mice were housed with free access to food and water in a temperature and humidity regulated room with a 12hr light/dark cycle.

The genotypes of the Par1b/MARK2-/- mice were determined with PCR using tail or ear genomic DNA with the following oligonucleotides: primer 1 MARK2-Common-F (5’- AGCCACCTCTGCTGACGAGCAGCC-3’), primer 2 MARK2-WT-R (5’-GCAGACTGAACCAGGACTTGGTGC-3’), and primer 3 MARK2-Mut-R (5’-ATGATCTGGACGAAGAGCATCAGG-3’). Primers 1 and 2 amplify a 310 bp fragment specific to the wild-type allele. Primers 1 and 3 amplify a 437 bp fragment specific to the mutant allele.

The genotypes of the Thy1-YFP mice were determined with PCR using the following oligonucleotides: primer 1 Transgene-R (5’- CGGTGGTGCAGATGAACTT-3’), primer 2 transgene-F (5’- ACAGACACACACCCAGGACA-3’), primer 3 Internal Positive Control-F (5’- CTAGGCCACAGAATTGAAAGATCT-3’), primer 4 Internal Positive Control-R (5’- GTAGGTGGAAATTCTAGCATCATCC-3’). Primers 1 and 2 amplify a 415 bp fragment specific to the transgene. Primers 3 and 4 amplify a 324 bp fragment specific to the internal positive control.

### Behavior

### Locomotor Activity

A standard plastic cage without bedding was placed in the Opto-Varimex Minor locomotor activity monitor. Locomotor behavior was recorded by the monitor based upon interruptions in horizontal and ambulatory photocell beam located throughout the cage. MARK2+/+ (n=14), MARK2+/- (n=12) and MARK2-/- (n=10) mice were individually housed for one week prior to the 30-minute test session.

### Rotarod Test

The rotarod test evaluates motor coordination and balance. A shortened latency for the mouse to fall from the rod indicates a deficit in motor coordination and balance. MARK2+/+ (n=14), MARK2+/- (n=12) and MARK2-/- (n=10) mice were placed on the rotarod with a diameter of 6 cm, rotating at 12 revolutions per minute. The rotarod was 60 cm above a padded receptacle. The latency to fall from the rotarod was recorded for each mouse for three trials, with each trial lasting 60 seconds.

### Elevated Plus Maze

The elevated plus maze (EPM) was used to measure anxiety-like behavior. More time spent in the open arms represents decreased anxiety-like behavior. The EPM was placed 60 cm above the floor with two long closed arms (65 cm long and 8 cm wide), two short open arms (30 cm long and 9 cm wide), and a central neutral 5 cm by 5 cm square. Each mouse was placed in the center square and observed for 10 minutes. The number and time the animal crossed into an open arm were recorded.

MARK2+/+ (n=13), MARK2+/- (n=12) and MARK2-/- (n=8) mice were tested.

### Morris Water Maze

The apparatus for the Morris Water Maze (MWM) consisted of a circular, steel tub (diameter: 110 cm; height: 59 cm), and for the purposes of behavioral scoring, was demarcated into four ‘imaginary’ quadrants (quadrant 1: Q1; quadrant 2: Q2; quadrant 3: Q3; quadrant 4: Q4). The tub was filled with regular tap water and water temperature was maintained at approximately 21 ± 1°C. The water was made opaque by adding black, non-toxic, powder tempera paint (Rich Art, Northvale, NJ). The platform consisted of a small, clear, perforated, plexiglass disc (diameter: 9 cm) mounted onto a steel rod and affixed to a heavy metal base (height: 48 cm). MARK2+/+ (n=12), MARK2+/- (n=14) and MARK2-/- (n=9) mice were tested for seven consecutive days, four trails per day. In each trial, a maximum swim time of 60 s was imposed. Each animal was subject to 2 days of visible platform training followed by 4 days of hidden platform test and a probe test on Day 7.

For the visible platform test (Day 1 and Day 2), the platform with a flag was raised 1 cm above the water. Animals were trained without the cues on the wall. The platform and start positions were randomly changed for each trial to avoid habituation to a particular quadrant.

For the hidden platform test (Day 3 through Day 6), the platform was submerged 1 cm below the surface of the water and placed in Q3. The start positions were randomly changed. Animals were trained with the cues on the wall. In each trial, a maximum swim time of 60 s was imposed. Between trials, a 10 s interval was imposed with the mouse on the platform. Spatial learning was measured as the time the mouse spent to find the platform. A shorter latency to find the platform represents better spatial learning.

For the probe test (Day 7), the platform was removed. During this probe trial the mouse was allowed to swim for 60 s. A significant increase in time spent in the target quadrant compared with all other quadrants represents retention of spatial memory.

### T-maze Active Avoidance Test

Mice were tested in a T-maze consisting of two 20 × 11 cm chambers connected to a 40 × 10 cm corridor with 18 cm high walls made of plexiglass. The floor was made of stainless-steel bars spaced 0.75 cm apart and connected to a shock generator except in the “safe” chamber. In each trial mice were placed in the start box with an initial interval of 20 s. A conditional stimulus tone accompanied by the opening of the start box door initiated the trial. Mice can avoid the shock by moving to the safe arm of the T-maze within 10 s of the tone. Failure to make an avoidance response led to onset of a 0.8 mA foot shock, which could be terminated by moving to the safe arm as an escape response (also 10 s maximum). Each animal underwent 10 trials per day, per mouse, per group (MARK2+/+ n=10, MARK2+/- n=10 and MARK2-/- n=8). The type of response (avoidance or escape) and the latency for the animal to make either avoidance or escape response were recorded.

### Social Chamber

The social chamber was a 40 cm × 40 cm × 36.6 cm plexiglass chamber with a stainless-steel grid floor. Within the chamber there were two cylinders 11 cm in diameter and 13 cm tall made of the same stainless-steel grid as the floor, located in opposite corners of the chamber. Each mouse was given a 10-minute habituation period to explore the chamber before the start of the trial. After 10 minutes one age- and sex-matched MARK2+/- mouse (target mouse) was placed in one of the cylinders and the subject mouse was placed in the center of the chamber. A contact was recorded each time the subject placed one or both paws on a cylinder. The number of contacts with either the target cylinder containing the target mouse or the control cylinder containing nothing was recorded. More contact with the target cylinder containing a novel mouse compared to the empty cylinder indicated a higher level of social behavior.

### Seizure Susceptibility

MARK2+/+ and MARK2+/- mice were injected with pilocarpine (300 mg/kg) intraperitoneally to induce seizures. To reduce side effects from peripheral cholinergic inputs, mice were also injected with methyl scopolamine (1 mg/kg) 30 minutes prior to pilocarpine injection. Mice were recorded for 2 hours following the injection and seizure behavior was scored by a blinded observer using the Racine scale as modified by Borges et al(53). Classifications were as follows: normal activity (stage 0); rigid posture or immobility (stage 1); stiffened, extended, and often arched tail (stage 2); partial body clonus, including forelimb or hind limb clonus or head bobbing (stage 3); whole body continuous clonic seizures with rearing (stage 4); severe whole body continuous clonic seizures with rearing and falling (stage 5); and tonic-clonic seizures with loss of posture or jumping (stage 6).

### Western Blot

For Western blot, whole brain tissue was lysed on ice in buffer containing 25 mM Hepes, 150 mM NaCl, 10 mM MgCl_2_, 1% Nonidet P-40, and 10 mM DTT and supplemented with protease inhibitor cocktail (1:1000, Sigma Aldrich), phosphatase inhibitor cocktail (1:100, Sigma Aldrich), 10 mM β- glycerophosphate, and 10 mM NaF. Lysates were cleared by centrifugation at 13,000×g for 10 min at 4 °C. Primary and secondary antibodies used are listed in **S1 Table**. Proteins were visualized by enhanced chemiluminescence and imaged using a Syngene G:BOX iChemi XR system (Syngene USA, Frederick, MD) or Azure 600 imaging system (Azure Biosystems Inc., Dublin, CA). Expression was quantified in FIJI ImageJ and normalized to GAPDH as a loading control.

### Dendritic Spine Analysis

MARK2 KO/YFP-H mice were perfused with cold PBS and 4% paraformaldehyde (PFA) followed by a post fixation in 4% PFA overnight in 4°C, and gradient sucrose solutions (10%, 20% and 30/%), each concentration for 24 hours in 4°C. The brains were then sectioned at 70μm using a microtome. Following washes in 1X PBS, sections were mounted using Vectashield with DAPI (Vector Laboratories, Burlingame, CA).

Fluorescence images were acquired using an Olympus FV1000 confocal microscope with a 60X water-immersion lens (NA 1.00, Olympus, Center Valley, PA). Z-stacks of dendritic spines were taken in 0.5um steps at 60X from the *strata radiatum* layer of the hippocampus. Images of dendritic spines were blinded to the experimenter analyzing the images, and the morphology and density were measured using Reconstruct(70). Spine length was defined as the length from the tip of the spine head to the point where the spine joins the dendrite. Spine width was defined as the maximum width of the spine head perpendicular to the long axis of the spine neck.

### Electrophysiology

Brain slice physiology was performed as described previously(71) with modifications. Briefly, mice were deeply anesthetized with Euthasol. Brains were removed and quickly immersed in cold (4°C) oxygenated cutting solution containing the following (in mM): 50 sucrose, 2.5 KCl, 0.625 CaCl_2_, 1.2 MgCl_2_, 1.25 NaH_2_PO_4_, 25 NaHCO_3_, and 2.5 glucose (oxygenated with 95%O_2_/5%CO_2_). Coronal hypothalamic or VTA slices, 300 μm in thickness, were cut using a vibratome (catalog #VT 1200S, Leica). Brain slices were collected in artificial CSF (ACSF) and bubbled with 5% CO_2_ and 95% O_2_. The ACSF contained the following (in mm): 125 NaCl, 2.5 KCl, 2.5 CaCl_2_, 1.2 MgCl_2_, 1.25 NaH_2_PO_4_, 26 NaHCO_3_, and 2.5 glucose, bubbled with 95%O_2_/5%CO_2_. After 60 min of recovery, slices were transferred to a recording chamber and perfused continuously at 2-4 ml/min with oxygenated ACSF at 30 °C. To record mini excitatory postsynaptic currents (mEPSCs), picrotoxin (50 µM, Sigma Aldrich, Saint Louis, MO) was added to block inhibitory currents mediated by GABA receptors and TTX (1 µM, Abcam Biochemicals) was added to block action potentials. Patch pipettes (3.8-4.4 MΩ) were pulled from borosilicate glass (G150TF-4; Warner Instruments) and filled with internal solution containing (in mM): 40 CsCl, 10 HEPES, 0.05EGTA, 1.8 NaCl, 3.5 KCl, 1.7 MgCl_2_, 2 Mg ATP, 0.4 Na_4_GTP, 10 Phosphocreatine (pH 7.2, 280-290 mOsm). EPSCs were recorded in whole-cell voltage clamp (Axon 100B amplifier, Molecular Devices), filtered at 2 kHz, digitized at 10 kHz, and collected using Clampex 10.2 (Molecular Devices). Neurons were voltage-clamped at -70 mV.

### RNA-seq

MARK2+/+ and MARK2-/- mice were sacrificed at 8 weeks old via live cervical dislocation followed by hippocampal dissection on ice cold PBS and flash freezing in liquid nitrogen or dry ice.

Hippocampal samples were stored at -80°C until they were sent to Admera Health (South Plainfield, NJ) for extraction, library preparation, and sequencing. Libraries were prepared with NEBNext UltraII (non- directional) kit with Poly A selection. Data was processed using 150-bp paired end reads. Adaptor sequences were removed from raw sequences using CutAdapt (72) and then remaining reads were aligned to the mm10 mouse genome (NCBI RefSeq Assembly GCF_000001635.20) using STAR v.2.7.10a (73). Reads with mapping quality below 30 were removed with Samtools (74). RNA-seq data analysis was performed with DESeq2 (75) in RStudio. Genes with low expression (less than 20 counts) were excluded. DESeq2 was also used to calculate TPM, FPKM, fold changes, P values, and adjusted P values (P_adj_). P_adj_ significance threshold was 0.05. Heatmaps (pheatmap (76)), MA (ggplot2), and volcano (ggplot2) plots were created in RStudio. Gene ontology (GO) analysis was performed using DAVID (77,78). Protein-protein-interaction networks were created with the STRING database using Cytoscape (79) and MCL clustering with AutoAnnotate (80).

### Statistical Analysis

All statistical analyses were performed using GraphPad Prism. All datasets were tested for normality. If not normal, ROUT (Q=1%) was used to normalize. Non-parametric tests were used if no outliers were found. Unpaired t-tests or Mann Whitney U tests were used to compare between two groups. Ordinary one-way ANOVA, Kruskall Wallis, or Brown Forsythe, were used for comparisons between three or more groups followed by Tukey’s *post hoc* test. Nested one-way ANOVAs were used in place of ordinary one-way ANOVA for spine analyses of length, width, length-to-width ratio (LWR), and spine density to account for individual mouse differences. Two-way ANOVA was used for spine morphology analysis followed by Tukey’s *post hoc*. Repeated measures two-way ANOVA was used for behavioral tests that spanned multiple time points followed by Dunnett’s *post hoc* test.

## Acknowledgements

This work was supported by NIH grant NS089578 and by the Assistant Secretary of Defense for Health Affairs endorsed by the Department of Defense, through the Epilepsy Research Program under Award No. W81XQH-18-1-0338 to HZ. Opinions, interpretations, conclusions, and recommendations are those of the author and are not necessarily endorsed by the Department of Defense. The U.S. Army Medical Research Acquisition Activity, 820 Chandler Street, Fort Detrick MD 21702-5014 is the awarding and administering acquisition office.

