## Supplemental Figures for "Neuronal connectivity, behavioral, and transcriptional alterations associated with the loss of MARK2"

* Corresponding author

¶ These authors contributed equally.

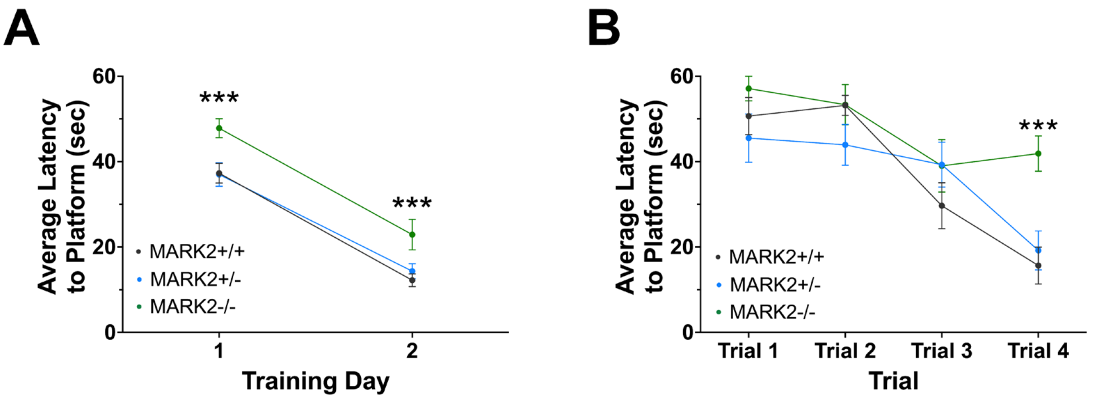


**S1 Fig. Morris Water Maze Days 1 and 2**

**(A)** Average latency to platform (in seconds) when platform was visible during training days 1 and 2. Two-way repeated measures ANOVA followed by Dunnett's *post hoc* test.

**(B)** Individual trials of training day 1. All mice successfully found the platform during the first three trials; however, MARK2-/- mice had increased latency by trial 4. Two-way repeated measures ANOVA followed by Dunnett's *post hoc* test.

*p<0.0332, **p<0.0021, ***P<0.0002, **** p<0.0001. Mean +/- SEM.

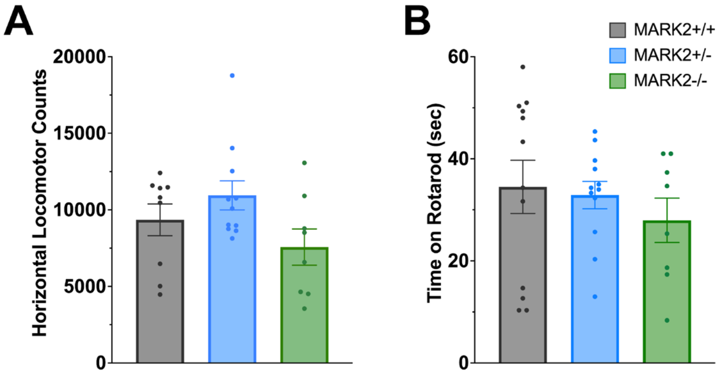


**S2 Fig. Locomotor and Rotarod**

**(A)** No changes were observed in locomotor activity across genotypes. Activity was measured in MARK2+/+ (n=14), MARK2+/- (n=12) and MARK2-/- (n=10) mice by quantifying the number of times they crossed the photocell beams in the cage. Kruskall-Wallis (stat=3.171, p=0.2048).

**(B)** MARK2+/+ (n=14), MARK2+/- (n=12) and MARK2-/- (n=10) showed no differences in performance on the rotarod test. Brown Forsythe ANOVA (F(2, 26.1)=0.3080, p=0.7375).

Mean +/- SEM plotted.

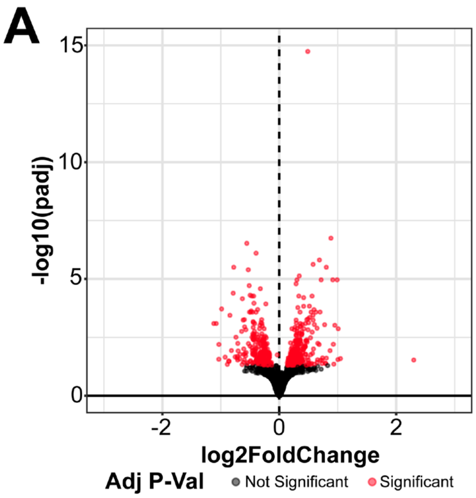


**S3 Fig. Volcano Plot of bulk RNAseq of MARK2+/+ and MARK2-/- hippocampi.**

**(A)** Volcano plot of 15,625 genes from DESeq2 analysis of 8-week-old MARK2+/+ (n=4) and MARK2-/- (n=5) mouse hippocampi. 522 significant genes are shown in red (Padj < 0.05).

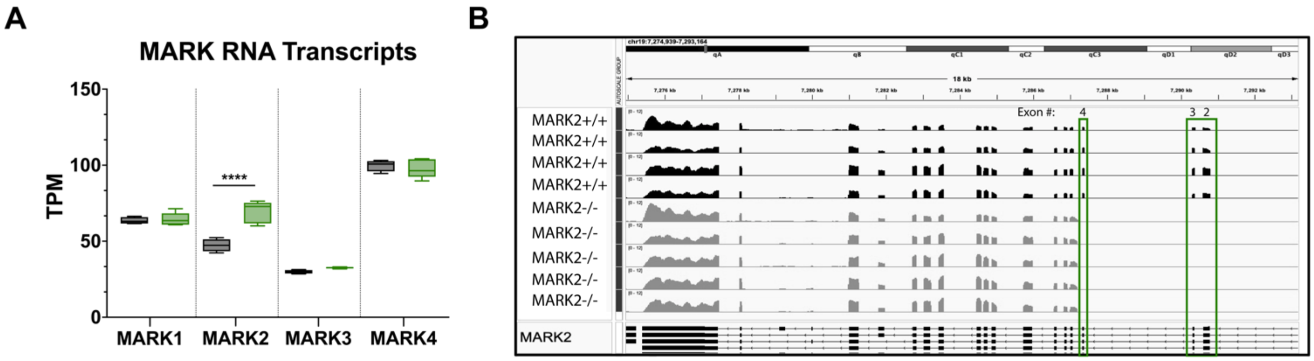


**S4 Fig. MARK family gene expression from RNAseq.**

**(A)** Transcripts per kilobase million (TPM) of MARK family genes in MARK2+/+ and MARK2-/- mice.

**(B)** Confirmation that exons 2-4 are absent from *MARK2* in the MARK2-/- mice.

**S1 Table. Antibodies used in WB experiments**

| Antibody | Concentration | Catalog # | Company | Application |
|---|---|---|---|---|
| RαMARK1 | 1:1000 | 21552-1-AP | ProteinTech | WB |
| RαMARK2 | 1:1000 | 15492-1-AP | ProteinTech | WB |
| MαMARK3 | 1:2000 | #05-680 | Millipore | WB |
| MαGAPDH | 1:8000 | #MAB374 | Sigma-Aldrich | WB |
| GαR HRP | 1:5000 | 111-035-144 | Jackson ImmunoResearch Laboratories | WB |
| GαM HRP | 1:5000 | 115-035-003 | Jackson ImmunoResearch Laboratories | WB |

### Additional Supplemental Files

S1 File. RNAseq DESeq2 Results.

S2 File. RNAseq TPM Results.

S2 File. DAVID Gene Ontology.

S3 File. Statistics Information.
